# The Cryo-Electron Microscopy Structure of the Type 1 Chaperone-Usher Pilus Rod

**DOI:** 10.1101/164657

**Authors:** Manuela K. Hospenthal, Tiago R. D. Costa, Adam Redzej, James Lillington, Gabriel Waksman

## Abstract

Chaperone-usher pili are long, polymeric protein fibres displayed on the surface of many bacterial pathogens. These critical virulence factors allow bacteria to specifically attach to host cells during infection. The type 1 and P pili of uropathogenic *Escherichia coli* (UPEC) play important roles during UPEC’s colonisation of the urinary tract, mediating bacterial attachment to the bladder and kidney, respectively. Also, their biomechanical properties that allow them to reversibly uncoil in response to flow-induced forces are critical for UPEC’s ability to retain a foothold in the unique and hostile environment of the urinary tract. Here we provide the 4.2 Å resolution cryo-electron microscopy (cryo-EM) structure of the type 1 pilus rod, which together with the previous structure of the P pilus rod, enables us to understand the remarkable “spring-like” properties of chaperone-usher pili in more detail.

## INTRODUCTION

Chaperone-usher pili are long, thin surface appendages displayed by many pathogenic Gram-negative bacteria (Thanassi et al., 1998). They serve to mediate important processes such as bacterial attachment to host tissues and biofilm formation, making them key virulence factors (Hospenthal et al., 2017; Thanassi et al., 2012). The two archetypal chaperone-usher pili are the type 1 and P pili of uropathogenic *Escherichia coli* (UPEC). UPEC are responsible for ~80% of community-acquired urinary tract infections (UTIs) and the role that chaperone-usher pili play in UTIs is firmly established (Flores-Mireles et al., 2015; McLellan and Hunstad, 2016).

The architecture of type 1 and P pili consists of a long, helically wound rod and a thin, flexible tip fibrillum located at the pilus’ distal end (Sauer et al., 2004) (Figure S1). The subunit located at the very tip of the fibrillum is the adhesin (FimH for type 1 and PapG for P pili). The adhesin consists of a C-terminal lectin domain responsible for host cell receptor interaction and an N-terminal pilin domain that links to the next subunit in assembly (Choudhury et al., 1999; Dodson et al., 2001). Two additional subunits, present as single copies, FimG and FimF (in that order; Figure S1), complete the structure of the type 1 pilus tip fibrillum. The P pilus tip fibrillum is slightly longer and consists of one subunit of PapF, 5-10 subunits of PapE, and one molecule of the adaptor subunit PapK. FimA (type 1 pilus) and PapA (P pilus) assemble into a ~1000 subunit long helically coiled quaternary structure of ~3-4 subunits per turn known as the rod (Habenstein et al., 2015; Hospenthal et al., 2017; 2016) (Figure S1).

To assemble chaperone-usher pili, all pilus subunits or “pilins” are first transported into the periplasm by the Sec YEG translocon (Stathopoulos et al., 2000) (Figure S1). There, they are folded and stabilised by a dedicated periplasmic chaperone, which shuttles each subunit to an outer membrane-embedded nanomachine termed the usher. On their own, pilins are unstable because they consist of an incomplete immunoglobulin (Ig)-like fold lacking the 7^th^ β-strand, which exposes a hydrophobic groove on their surface (Barnhart et al., 2000; Choudhury et al., 1999; Sauer et al., 1999; Vetsch et al., 2004). This groove can be complemented by a donor-strand originating either from the periplasmic chaperone in a process termed “donor-strand complementation (DSC)”, or from the next subunit in assembly in a process termed “donor-strand exchange (DSE)” (Nishiyama et al., 2008; Sauer et al., 2002; Zavialov et al., 2003). A region consisting of 10-20 N-terminal residues known as the N-terminal extension (Nte), present on all subunits except for the adhesin, serves as the donor-strand during DSE (Sauer et al., 2002; Waksman and Hultgren, 2009; Zavialov et al., 2003).

The usher receives chaperone-subunit complexes, polymerises the subunits into a pilus, and secretes the growing pilus to expose it on the bacterial surface (Geibel et al., 2013; Hospenthal et al., 2017; Phan et al., 2011; Remaut et al., 2008). The cycle of subunit incorporation by the usher is well-characterised (reviewed in Waksman, 2017): all chaperone-subunit complexes are first recruited to the N-terminal domain (NTD) of the usher and subsequently transferred to two C-terminal domains (CTDs) that form a secondary chaperone-subunit binding platform. DSE occurs when a subunit located at NTD reacts with the previously assembled subunit located at the CTDs. Indeed, the subunit at the NTD is positioned relative to the subunit at the CTDs in such a way that its Nte is close to the groove of the CTDs-bound subunit and thus can “zip-in” into that groove, thereby displacing the chaperone and forming a native Nte-groove subunit-subunit interaction. At this point, the NTD-bound chaperone-subunit complex transfers to the CTDs with a rotation and translation motion that results in the extrusion of the pilus, one subunit at a time.

Type 1 and P pilus rods exhibit remarkable biomechanical properties enabling UPEC to resist being flushed out of the urinary tract during a UTI. In response to urine flow-induced forces, the helically-coiled rod section of chaperone-usher pili can reversibly uncoil, thereby dissipating and relieving the force experienced by the adhesin-receptor complexes (Forero et al., 2006; Miller et al., 2006; Zakrisson et al., 2012). Several studies applying techniques such as atomic force microscopy (AFM) and optical tweezers have carefully deciphered and unpicked the processes that occur during the unwinding of the helical pilus rod (Andersson et al., 2008; 2006a; 2006b; 2006c; 2007; Fällman et al., 2005; Forero et al., 2006; Jass et al., 2004; Lugmaier et al., 2007; Miller et al., 2006; Zakrisson et al., 2012; 2013) and a mathematical model of the force vs elongation behaviour of pili was developed (Andersson et al., 2006a; Jass et al., 2004). Despite their similarities, type 1 and P pili behave differently in response to external forces. Notably, type 1 pili can respond faster to external force by entering a dynamic regime of elongation at lower elongation rates compared to P pili (6 nm/s vs 400 nm/s) (Andersson et al., 2007). In addition, type 1 pili require larger forces to peel apart the individual stack-to-stack interactions in the dynamic elongation mode (Andersson et al., 2008). It has been suggested that these differences allow type 1 pili to withstand the faster and more turbulent flows of the lower urinary tract, whereas P pili are biomechanically better evolved to allow colonisation of the upper urinary tract (Andersson et al., 2008).

The previously reported structure of the P pilus rod provided the molecular explanation of how the main stacking interface in the rod (formed by every n and n+3 subunit) can break apart, while the much stronger DSE forces holding adjacent subunits together remain intact (Hospenthal et al., 2016). Here we present the 4.2 Å resolution cryo-electron microscopy (cryo-EM) structure of the related type 1 pilus rod. The comparison of the type 1 and P pilus rod structures begins to explain some of the differences in their biomechanical properties at the molecular level.

## RESULTS

### Structure Determination and Architecture of Type 1 Pilus Rods

Type 1 pili were expressed and assembled on the *Escherichia coli (E. coli)* cell surface, from which they were sheared and purified by density gradient centrifugation as described in Experimental Procedures. The purified sample (Figure 1a) was applied to grids and vitrified for cryo-EM analysis (Figure 1b). The resulting electron density map was resolved to an overall resolution of ~4.2 Å (Figure S2a), which is consistent with the map showing clearly separated strands and visible density for bulky side chains (Figure 1c, d). An analysis of the local resolution of the EM map shows that the interior of the pilus rod is better resolved than the exterior (Figure S2b) and the lowest resolution is observed in outward-facing loops (including residues 66-67, 93- 95, 108-109, 116-117 and 140-143), suggesting some disorder in these regions. Nevertheless, a near-atomic resolution model of the fully assembled type 1 pilus rod was built by fitting a previously determined NMR solution structure of FimA (Puorger et al., 2011) into the EM map, followed by manual building and refinement (Figure S2c).

**Figure 1:**
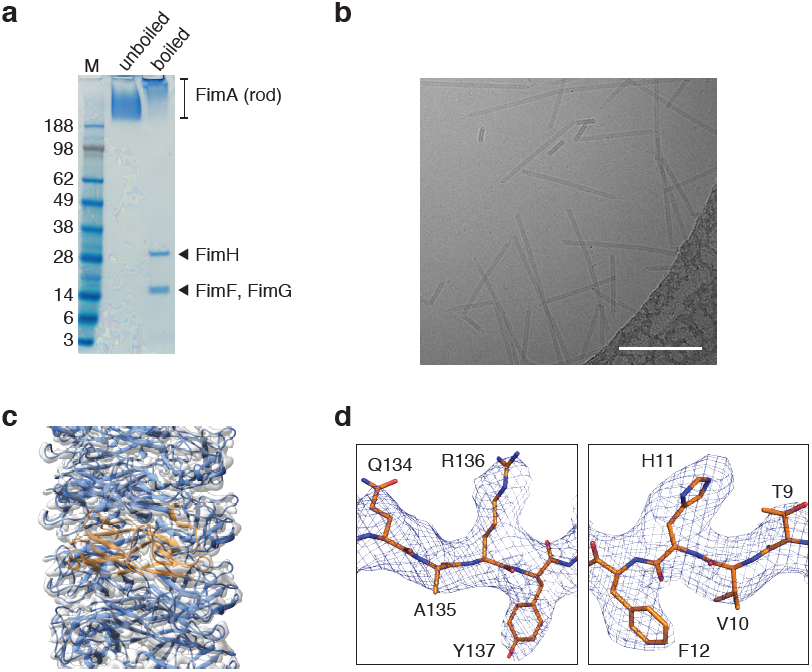
Purification and Cryo-EM Analysis of Type 1 Pili. **a** SDS-PAGE analysis of type 1 pili purified from the *E. coli* cell surface. When the sample was boiled for 15 min in the presence of SDS and 4.5 M urea, the pili partially dissociated and bands representing the monomers of some type 1 pilins were resolved. Mass spectrometry (LC-MS/MS) was performed to confirm the identity of all the bands (labelled). **b** A representative electron micrograph of type 1 pili. The scale bar represents 100 nm. **c** The refined type 1 pilus rod model is shown in ribbon representation in the experimentally derived electron density map (transparent grey surface). FimA subunits are coloured blue, with the central FimA subunit highlighted in orange. **d** Regions of electron density around bulky side chains. Electron density is shown as a blue mesh and the model is shown in stick representation with carbons coloured in orange, oxygens in red and nitrogens in blue. Residues are clearly labelled.

The pilus rod is ~72 Å in diameter and contains a hollow central lumen, which is ~14 Å wide (Figure 2a). Overall, the type 1 pilus is a right-handed superhelical assembly comprising 3.13 subunits per turn with an axial rise of 8 Å per subunit and a pitch of 25 Å (Figure 2b). An extensive subunit-subunit interaction network maintains the integrity of the superhelical quaternary structure, where each subunit interacts with a total of eight other subunits, four preceding and four succeeding subunits (n interacts with +1, +2, +3 +4 and −1, −2, −3, −4). This is similar to what has been observed for the related P pilus structure, where each subunit interacts with a total of ten subunits (n also interacts with +5 and −5) (Hospenthal et al., 2016) (see below). By far the largest contribution to the subunit-subunit interaction network is made by the main stacking interface created by every n and n+3 subunits (Figure 2c and S3). The total buried surface area created by the interaction of the donor-strand complemented n and n+3 pilins is 1616.2 Å^2^ (or 1430.2 Å^2^ when the contribution of the donor-strand is not taken into account) (Figure S3b).

**Figure 2:**
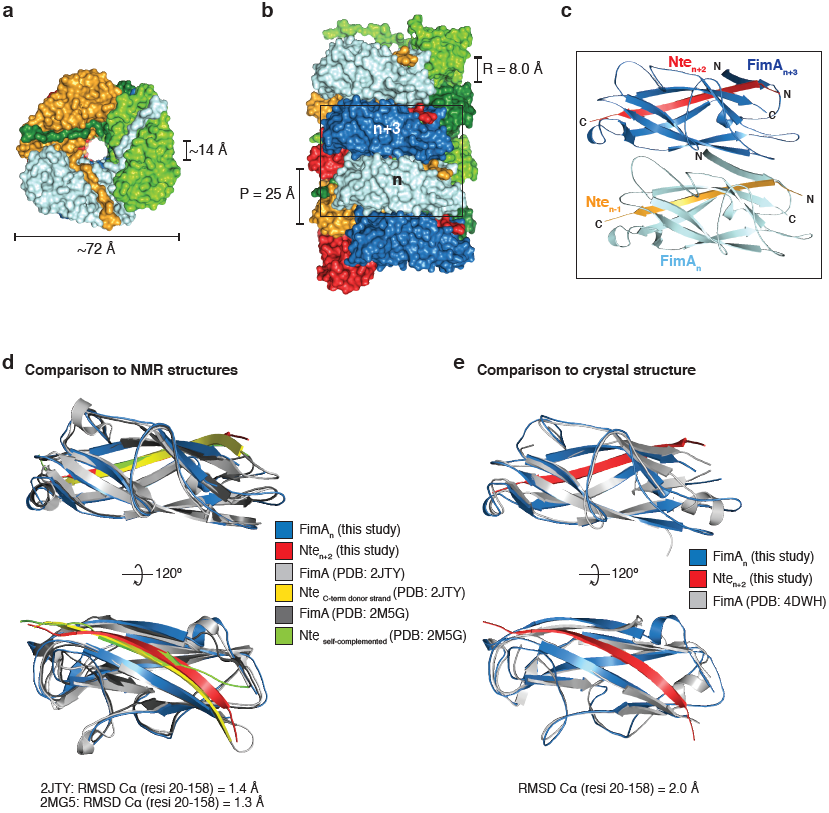
Overall Architecture of Type 1 Pili and Comparison of the FimA Pilin Structure. **a** Top view of the type 1 pilus rod model in surface representation. The last Nte of the light green molecule has been removed for clarity. The dimensions of the overall and lumen diameters are indicated. **b** Side view of the type 1 pilus rod model. There are three faces of the pilus rod structure, coloured in dark and light blue (front), dark and light green, and red and orange. A “stacking” interface is created between every n and n+3 subunit (boxed and labelled). The rise (R) and pitch (P) are indicated. **c** Zoomed-in view of the stacking interface boxed in b. The two FimA subunits are coloured in light and dark blue and are shown in ribbon representation. Donor-strands are coloured orange and red. Residue-specific interaction details for this interface are shown in Figure S3. **d, e** Superposition of the FimA pilin from the fully assembled pilus structure (dark blue) compared to **d** two NMR solution structures of FimA (PDB ID: 2JTY (Puorger et al., 2011), light grey; and 2M5G (Walczak et al., 2014), dark grey) and **e** a crystal structure of FimA (PDB ID: 4DWH (Crespo et al., 2012), light grey) shown in two views rotated 120° with respect to each other. All proteins are shown in ribbon representation and the RMSD values for the alignment of Cα atoms for residues 2-158 are indicated.

### FimA before and after Assembly into Type 1 Pili

Several structures of FimA have been determined before their assembly into pili. NMR was used to determine the structure of FimA in isolation where the hydrophobic groove was either self-complemented by FimA’s own Nte (FimA, PDB ID: 2M5G (Walczak et al., 2014)) or by a copy of the FimA Nte peptide which was fused to the C-terminus of the construct (FimAa, PDB ID: 2JTY (Puorger et al., 2011)). The two structures differ in the orientation of donor-strand insertion with respect to the last β-strand of the Ig-like pilin fold. In FimAa (PDB ID: 2JTY), the donor-strand is inserted in a stable antiparallel fashion, which mimics the orientation observed in the quaternary structure of the pilus; whereas in FimA (PDB ID: 2M5G) the donor-stand is inserted in a less stable parallel arrangement (Figure 2d). In addition, two structures of FimA interacting with the chaperone FimC were determined by X-ray crystallography (Crespo et al., 2012), but only the highest resolution structure (2.5 Å) (PDB ID: 4DWH) will be compared (Figure 2e). All three previous FimA structures were aligned to the structure of FimA assembled into a pilus using the pairwise alignment function of the DALI server (Holm and Laakso, 2016) (Figure 2d, e). The two solution structures of FimA align with a root-mean-square deviation (RMSD) (Cα) of 1.3 Å (FimA, PDB ID: 2M5G) and 1.4 Å (FimAa, PDB ID: 2JTY) for residues 20-158 (excluding the donor-strand), compared to an RMSD of 2.0 Å for the crystal structure (PDB ID: 4DWH), suggesting that the structures previously observed in solution are more representative of the structure of FimA in the context of the fully assembled pilus (Figure 2d, e). The main difference lies in a region of α-helix (residues 62-68) present in the crystal structure, which is absent and forms a loop in the structure of FimA inside the pilus (Figure 2e). In addition, there is a short 3_10_- helix (residues 25-28) present in FimA inside the pilus, which is absent in the crystal structure. The two NMR structures are very similar to FimA inside the pilus across residues 20-158, with perhaps the only difference being an additional short helix spanning residues 79-83 in FimAa (PDB ID: 2JTY), which is a loop in both the self-complemented FimA (PDB ID: 2M5G) and FimA inside the pilus (Figure 4d).

**Figure 4:**
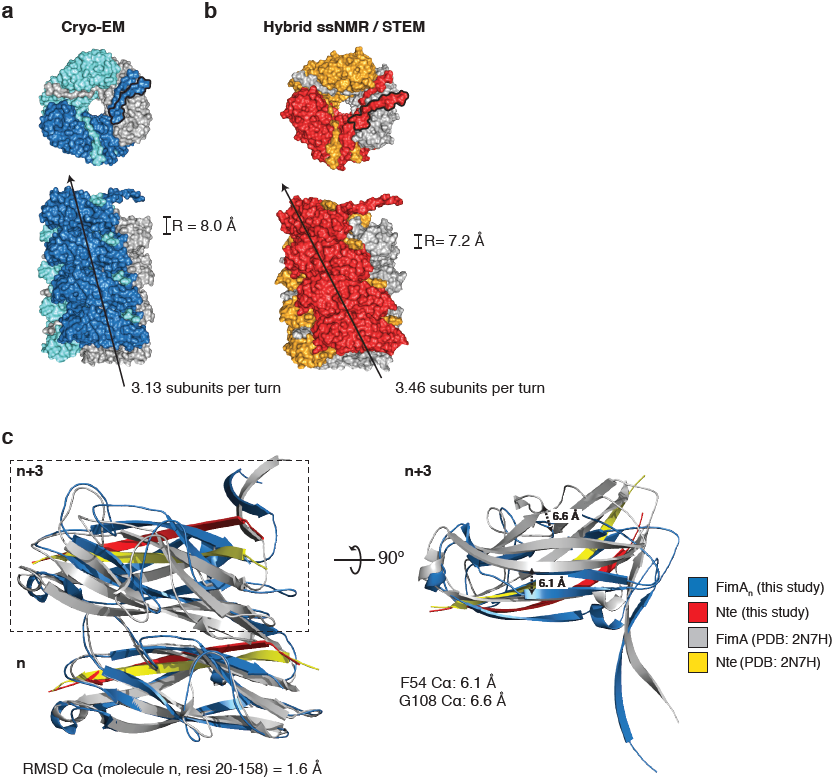
Comparison of the Type 1 pilus Rod Structure Determined by Cryo-EM and a Hybrid ssNMR / STEM Approach. **a** A top (upper panel) and side view (lower panel) of the cryo-EM derived model of the type 1 pilus rod shown in surface representation. The pilus subunits are coloured as in Figure 3a. A black arrow indicates the degree of twist in the pilus by tracing up the front face of the pilus structure. The helical parameter of rise (R) and the number of subunits per turn are indicated. **b** The hybrid model derived from ssNMR and STEM data (coloured in red, orange and grey) shown in surface representation and labelled as in a. The number of subunits per turn was calculated by dividing the pitch (24.9 Å) by the rise (7.2 Å) (Habenstein et al., 2015). The last Nte of the top subunit in the surface representation of the top view, which doesn’t DSE with another subunit, is outlined in black to distinguish it from the Nte of the same colour emanating from the subunit below (n-3). **c** Left panel, a superposition of the two subunits participating in the pilus’ main stacking interface (n and n+3) of the cryo-EM derived model (blue, Nte in red) and the ssNMR / STEM derived model (grey, Nte in yellow). The n subunit was aligned and RMSD value for the alignment of Cα atoms for residues 2-158 is indicated below. Right panel, a 90° rotation of the n+3 subunit showing the offset of key β-strands and loops in the stacking interface as a result of the differences in twist and rise. The distances between equivalent Cα atoms (F54 and G108) are indicated.

### Comparison of the Type 1 and P Pilus Rods

Comparing and contrasting the near-atomic resolution models of the type 1 and P pili provides a complete picture of the general architecture of the archetypal chaperone-usher pili from UPEC. The pilins are arranged into similar right-handed superhelical quaternary assemblies, which differ in their helical rise and the number of subunits required to complete one turn (Figure 3a, b). The P and type 1 pilus structures have a similar pitch but contain 3.28 and 3.13 subunits per turn respectively, making the P pilus slightly more tightly wound. This is also reflected in the slightly smaller helical rise of the P pilus structure (7.7 Å) compared to the type 1 pilus (8.0 Å) (Figure 3a, b). Both structures have an ascending path of Ntes that have their N-terminal ends facing the pilus exterior and their C-terminal ends lining the pilus lumen. The Nte of PapA is longer by one residue and its N-terminal portion (A1-P5), the so-called staple, creates a sharp ~90° angle with the remainder of the Nte (Hospenthal et al., 2016) (Figure 3c). By contrast, the N-terminal portion of the FimA Nte lies flat against the remainder of the FimA pilin fold. This difference is clear when the electron density of both the FimA and PapA Nte is compared, which allows the main chain to be traced unambiguously in both structures (Figure 3c).

**Figure 3:**
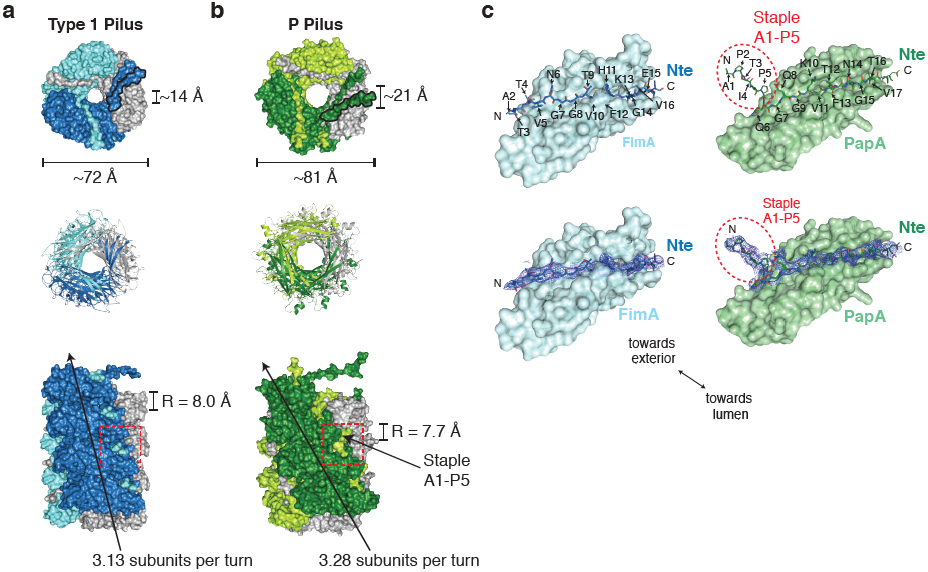
Comparison of the Type 1 and P Pilus Rod Structures. **a, b** Top and side views of **a** the type 1 pilus rod (coloured in dark blue, light blue and grey) and **b** the P pilus rod (coloured in dark green, light green and grey). The top view is shown in both surface and cartoon representation, whereas the side view is shown in surface representation. The last Nte of the top subunit in the surface representation of the top view, which doesn’t DSE with another subunit, is outlined in black to distinguish it from the Nte of the same colour emanating from the subunit below (n-3). The dimensions of the outer and lumen diameters (top view) and helical parameters of rise (R) and the number of subunits per turn are indicated (side view). A black arrow indicates the degree of twist in the pilus by tracing up the front face of the pilus structure. The N-terminal end of the Nte is visible between subunits (dashed red box) and is shorter and oriented differently in the type 1 pilus compared to the previously described “staple” region (residues 1-5) in the P pilus (Hospenthal et al., 2016). The model of the type 1 pilus begins at residue A2. **c** A comparison between the Nte peptides complementing the pilin’s hydrophobic groove in the type 1 pilus (left) and P pilus (right). The pilin subunits are shown in surface representation (FimA, light blue; PapA light green) and the Nte peptide is shown in stick representation (FimA, dark blue; PapA, dark green). The Ntes of the pilin subunits shown in surface representation have been removed for clarity. The bottom panels show the quality of the electron density surrounding the Nte peptides, illustrating the differences at the N-terminus. The PapA Nte makes a sharp turn and forms the “staple” region (red dashed circle), whereas the FimA Nte lies flat against the FimA subunit. Residues are labelled and an arrow indicates the overall orientation of the pilins.

### Comparison of Cryo-EM and ssNMR / STEM Structures of the Type 1 Pilus Rod

Interestingly, a model of the type 1 pilus rod has been determined using a hybrid approach of solid-state NMR spectroscopy (ssNMR) and scanning transmission electron microscopy (STEM) (Habenstein et al., 2015) (Figure 4). This model has a helical arrangement of 3.46 FimA subunits per turn with a rise of 7.2 Å (Figure 4b), which agrees with a previous low resolution EM study (Hahn et al., 2002). Although, our refined rise value of 8.01 Å falls within the range calculated using STEM mass-per-length (MPL) measurements (corresponding to 1.99 kDa/Å, in a range of 1.8-2.5 kDa/Å), such a discrepancy leads to significant offsets observed in the stacking interface between the n and n+3 subunits (Figure 4c). Furthermore, while the ssNMR / STEM study suggests that the sample of type 1 pili are fairly heterogeneous with respect to their helical parameters (rise range of 6.38-8.87 Å), our cryo-EM analysis points to a homogenous sample where the helical parameters do not vary greatly (Experimental Procedures and Table S1).

## DISCUSSION

This near-atomic resolution model of the type 1 pilus rod allows us to fully appreciate the similarities and differences of the two archetypal chaperone-usher pili of UPEC. FimA and PapA are related proteins with 31.7% sequence identity (mature proteins), sharing the same C-terminally truncated Ig-like pilin fold and are assembled by related members of the chaperone-usher pathway. Not surprisingly, the quaternary structure formed by FimA or PapA, the pilus rod, also shares several similarities. Both the type 1 and P pilus rods form right-handed superhelical arrangements of similar overall dimensions (Figure 3a, b). The P pilus rod is formed of 3.28 subunits per turn with an axial rise of 7.7 Å, whereas the type 1 pilus rod is formed of 3.13 subunits per turn with a rise of 8.0 Å. These differences mean that the P pilus adopts a more “twisted” or tightly wound conformation (Figure 3b). Furthermore, the type 1 and P pilus rod structures differ in their Nte regions. The five N-terminal residues (A1-P5) of PapA form the “staple” region which forms a ~90° angle with respect to the remainder of the Nte (Figure 3c). This difference in the conformation (and length) of the Nte allows the staple residues of PapA to reach and make contact with the n+5 subunit, thereby creating an interaction network where each subunit contacts ten others (two more than in the type 1 pilus) (Hospenthal et al., 2016).

However, as demonstrated for the P pilus, the most important interface is the main stacking interface formed between every n and n+3 subunit (Figure 2c and S3a). This interface is responsible for maintaining the quaternary structural integrity of the pilus and also governs the biomechanical properties of reversible uncoiling in response to shear forces such as those experienced in the urinary tract. Chaperone-usher pili have been the subjects of several studies utilising force spectroscopy techniques such as optical tweezers or atomic force microscopy (AFM) (Andersson et al., 2008; 2007; 2006c; 2006b; 2006a; Fällman et al., 2005; Forero et al., 2006; Jass et al., 2004; Lugmaier et al., 2007; Miller et al., 2006; Zakrisson et al., 2013; 2012). A mathematical model of the force vs elongation behaviour of P pili was developed, identifying three elongation regions (Jass et al., 2004). Region I is characterised by a linear force vs elongation response and is thought to reflect the elastic stretching of the quaternary rod structure (although not yet breaking it). Region II results in elongation under constant force and represents the sequential opening of the stack-to-stack interactions resulting in rod unwinding. Finally, Region III shows an “s-shaped” force vs elongation response and represents the overstretching of the now linearised rod, still held together by the donor-strand exchange interactions. Both regions I and II depend on the interface created by the n and n+3 subunits. The unwinding of the rod in Region II occurs either under steady-state or dynamic conditions depending on the elongation speed applied (Andersson et al., 2008; 2007). Measurements performed under steady-state conditions, revealed that type 1 and P pili unwind at comparable unfolding forces (28 ± 2 and 30 ± 2 pN, respectively) (Andersson et al., 2008). However, measurements performed under dynamic conditions (dynamic force spectroscopy (DFS)) can address values of physical entities that steady-state measurements cannot address, such as the bond-opening rate and bond lengths (bond here refers to the stack-to-stack interactions in the quaternary rod structure) (Andersson et al., 2008). Such measurements have suggested that a higher force is required to unwind type 1 pili compared to P pili at these fast elongation rates. In turn, this implies that the stacking interface is stronger in type 1 pili (Andersson et al., 2008). These findings are indeed supported by our cryo-EM structure of the type 1 pilus rod, which has a larger buried surface area (1616.2 Å^2^) in the n and n+3 (stacking) interface than the P pilus (1453.0 Å^2^) (Hospenthal et al., 2016) (Figure S3b). Given that the two interfaces have a similar composition in terms of polar and non-polar residues, this larger buried surface area may explain the lower thermal bond opening rate observed for type 1 pilus rods.

Interestingly, two different recoiling forces have been observed for type 1 pili, suggesting that type 1 pili can rearrange into two distinct quaternary configurations after having been extended and linearised (Andersson et al., 2008; 2007). This raises the intriguing question of whether two different structures of type 1 pili, the cryo-EM structure (this study) and the hybrid ssNMR / STEM model (Habenstein et al., 2015), may represent the two different states observed in these force spectroscopy experiments. While our type 1 pilus sample was purified from the surface of *E. coli* (see Experimental Protocols) and resulted in analysed segments that were homogeneous with respect to the helical twist and rise (Table S1), the sample used for the ssNMR / STEM measurements was first unfolded and then refolded *in vitro*. Therefore, it is conceivable that the cryo-EM structure represents pilus rods after assembly by the usher, whereas the ssNMR / STEM structure represents a structure that followed a different refolding path. The range of values observed in the MPL experiment suggests that these filaments had more than a single quaternary conformation. In addition, factors such as pH and salinity can affect the refolding / recoiling behaviour of pili during force spectroscopy measurements (Andersson et al., 2008), which raises the additional possibility that low pH stains (e.g. uranyl acetate or uranyl formate) could affect the quaternary structure of pilus rods. Indeed, previous studies used uranyl formate (pH 4.25) stained grids to determine the helical parameters of type 1 pili (Hahn et al., 2002). Whether or not such factors can affect the helical parameters of type 1 pili and whether such different structures do indeed exist, remains to be fully determined.

Chaperone-usher pili are biopolymers with remarkable biomechanical properties. The availability of near-atomic resolution models of both the type 1 and P pilus rods provides an unprecedented opportunity to understand what factors govern their differences when it comes to reversible rod uncoiling. Although these structures appear superficially similar, subtle differences in key interfaces and in their helical parameters could influence how these structures behave in response to shear forces. The work described here, together with that of the P pilus (Hospenthal et al., 2016), now provides the basis on which accurate molecular dynamics simulations can be implemented, that will probe the energetic, structural and molecular basis of pilus uncoiling / recoiling.

## EXPERIMENTAL PROCEDURES

### Plasmids and Bacterial Strains

The pSH2 plasmid harbouring the *fim* operon from the UPEC strain J96 has been described previously (Orndorff and Falkow, 1984). An arabinose-inducible plasmid, pSH5, was created by subcloning the *FimA* to *FimH* gene cluster from pSH2 onto the pBADM-11 vector backbone. pSH5 was transformed into HB101 *E. coli* cells (Promega) for cell-surface pilus production.

### Type 1 Pilus Expression and Purification

Cultures of HB101 *E. coli* cells, transformed with pSH5, were grown to OD_600_ of 0.6-0.8 in Luria-Bertani (LB) media at 37°C in a shaking incubator. The expression of type 1 chaperone-usher pili was induced with 0.05% (w/v) arabinose and the culture was incubated further for 1 h at 37°C before harvesting the cells by centrifugation. Pili were sheared off the cell surface by gently stirring the cells for 2 h resuspended in 400 mL of buffered solution containing 30 mM sodium citrate dehydrate [pH 7.2], 300 mM NaCl, 1 mg/mL DNase and complete mini EDTA-free protease inhibitor cocktail tablets (Roche). Subsequently, depiliated cells were removed by two rounds of centrifugation at 10800 *g* for 30 min and pili were precipitated by incubating and gently stirring the supernatant in the presence of 5% (w/v) PEG 6000, 0.5 M NaCl for 30 min. The precipitate was pelleted by centrifugation at 18000 *g* for 30 min. This pellet was resuspended in 60 mL of Milli-Q water and gently stirred for 20 min at 22°C before centrifugation at 5000 *g* for 20 min. The supernatant containing pili was precipitated once more in the presence of 5% (w/v) PEG 6000, 0.5 M NaCl as previously before a final centrifugation step at 27000 *g* for 30 min. Pili were resuspended in 500 μL of buffer A (20 mM Tris [pH 7.5], 150 mM NaCl) before being layered onto a pre-formed CsCl step gradient (1.1-1.4 g/cm^3^) and centrifuged at 260000 *g* for 17 h. The pili containing band was carefully removed and dialysed against buffer A. Once fully dialysed, the pili containing solution was applied to a 15%-60% (w/v) sucrose density gradient and centrifuged at 160000 *g* for 12 h. Fractions from the sucrose density gradient were analysed by SDS-PAGE using 4%-12% NuPage gradient gels (Life Technologies) and pili containing fractions were pooled. The final sample was dialysed against buffer A and the identity of type 1 chaperone-usher pili was confirmed by the identification of FimA, FimF, FimG and FimH by mass spectrometry. All purification and centrifugation steps were carried out at 4°C unless stated otherwise.

### Cryo-Electron Microscopy Sample Preparation and Data Collection

A 3 μL sample of type 1 chaperone-usher pili was applied to a glow-discharged Quantifoil 1.2/1.3 400 mesh grid (Agar Scientific) and incubated for 30 s before being blotted and plunged into liquid ethane using a Vitrobot plunge-freezing device (FEI). The data were collected on a Tecnai G^2^ Polara microscope (FEI) operated at 300 kV equipped with K2 Summit direct electron detector (Gatan) with a 1.13 Å pixel size and a defocus range of −0.5 to −3.5 μm. A total dose of 100 electrons/Å^2^ was applied and fractionated equally among 59 frames to allow for dose weighting.

### Cryo-Electron Microscopy Image Processing and Reconstruction

Whole-image drift correction to align the 59 movie frames of each micrograph was carried out using MOTIONCOR2 (Zheng et al., 2017) and the contrast transfer function (CTF) parameters of the corrected micrographs were estimated using GCTF (Zhang, 2016). The implementation for the reconstruction of helical assemblies in the program RELION-2.0 (He and Scheres, 2017; Scheres, 2012) was used for image processing and reconstruction. Filaments were manually picked from 177 selected micrographs and a total of 115545 segments were extracted with a box size of 240 pixels. After 2D and 3D classification steps, a total of 115510 segments were used for 3D refinement. No segments were discarded after 3D classification, as all three classes refined with highly similar helical parameters. A solid cylinder with a diameter of 100 Å, low-pass filtered to 30 Å, was used as the starting model for 3D reconstruction and the helical parameters of twist and rise were refined from an initial narrow search range. During the post-processing step in RELION-2.0, a soft mask with a raised cosine edge 7 pixels wide was employed yielding a final map with a global resolution of 4.2 Å as assessed by the gold standard FSC procedure implemented in RELION-2.0 (FSC=0.143) (Rosenthal and Henderson, 2003), consistent with strand separation and clear density for bulky side chains.

### Model Building and Refinement and Structure Analysis

The FimA pilin (PDB ID: 2JTY (Puorger et al., 2011)) structure encompassing residues 20-159 was docked into the map using Chimera. The donor-strand was initially modelled with residues 172-182 (representing Nte residues 7-17) from the self-complemented donor strand of the same FimA pilin structure (PDB ID: 2JTY). The Nte residues were renumbered and missing residues (2- 6 and 18-19) were manually built using *Coot* (Emsley et al., 2010). The final model encompasses residues 2-158 of FimA, missing one residue from both the N- and C-terminus due to poorly defined electron density. A model containing six molecules of FimA was refined using several cycles of PHENIX (Real Space Refine) (Adams et al., 2010). Manual adjustment in *Coot* and structure idealisation in REFMAC5 (Vagin et al., 2004) was performed to improve the geometry of the model between cycles of real space refinement. Knowledge of the FimA structure from previous NMR and crystallography studies were used to guide building and refinement (Crespo et al., 2012; Puorger et al., 2011; Walczak et al., 2014). However, in order to obtain an unbiased view of secondary structure element boundaries, the DSSP server (Joosten et al., 2010; Kabsch and Sander, 1983) was used in combination with careful manual examination of the model in *Coot* to delineate the final secondary structure element boundaries enforced during real space refinement. Final validation of the model was performed using MOLPROBITY (Chen et al., 2009) and the wwPDB validation Service. The pairwise alignment function of the DALI server was used to calculate RMSD values for the alignment of two structures across a range of Cα atoms (Holm and Laakso, 2016). The CoCoMaps Tool was used to analyse the interfaces between FimA or PapA pilins in the quaternary structure of their respective pili (Vangone et al., 2011).

## AUTHOR CONTRIBUTION

M.K.H. purified pili, prepared cryo-EM grids, collected data, performed image processing and reconstruction, model building, made figures and wrote the paper. T.R.D.C. and A.R helped with screening conditions for and vitrifying of cryo-EM grids and T.R.D.C. supervised EM data collection. J. L. cloned the pSH5 plasmid. G.W. supervised the work and wrote the paper.

### ACKNOWLEDGEMENTS

This work was funded by MRC grant 018434 to G.W. We wish to thank Dr Natasha Lukoyanova for assistance with EM data collection and Shaoda He for advice on image processing in RELION-2.0. We also wish to thank Dr Aravindan Ilangovan and Germán Sgro for advice on model building and refinement.

